# emiRIT: A text-mining based resource for microRNA information

**DOI:** 10.1101/2020.11.05.370593

**Authors:** Debarati Roychowdhury, Samir Gupta, Xihan Qin, Cecilia N. Arighi, K. Vijay-Shanker

## Abstract

**Motivation:** microRNAs (miRNAs) are essential gene regulators and their dysregulation often leads to diseases. Easy access to miRNA information is crucial for interpreting generated experimental data, connecting facts across publications, and developing new hypotheses built on previous knowledge. Here, we present emiRIT, a text mining-based resource, which presents miRNA information mined from the literature through a user-friendly interface.

**Results:** We collected 149,233 miRNA-PubMed ID pairs from Medline between January 1997 to May 2020. emiRIT currently contains *miRNA-gene* regulation (60,491 relations); *miRNA-disease (cancer)* (12,300 relations); *miRNA-biological process and pathways* (23,390 relations); and circulatory *miRNAs in extracellular locations* (3,782 relations). Biological entities and their relation to miRNAs were extracted from Medline abstracts using publicly available and in-house developed text mining tools, and the entities were normalized to facilitate querying and integration. We built a database and an interface to store and access the integrated data, respectively.

**Conclusion:** We provide an up-to-date and user-friendly resource to facilitate access to comprehensive miRNA information from the literature on a large-scale, enabling users to navigate through different roles of miRNA and examine them in a context specific to their information needs. To assess our resource’s information coverage, in the absence of gold standards, we have conducted two case studies focusing on the target and differential expression information of miRNAs in the context of diseases. Database URL: https://research.bioinformatics.udel.edu/emirit/

## 1. Introduction

MicroRNAs (miRNAs) are non-coding small RNAs that regulate gene expression at the post-transcriptional level. The majority of protein-coding genes are controlled by miRNAs, suggesting that most biological processes are subjected to miRNA-dependent regulation (1). Several studies have also shown miRNAs’ implications in cancer and neurodegenerative diseases (2–8). Experimental findings regarding miRNAs, covering various contents such as target information, differential expression of miRNAs, and their role in diseases, are scattered across multiple publications and databases (9, 10). For example, consider a biomedical researcher interested in knowing the miRNAs that are differentially expressed in the context of Triple-negative breast cancer. The researcher may be interested in the biological processes impacted by such miRNAs or their target genes, specifically if they are mentioned in this disease’s context. Such information may be found in multiple publications and different databases. However, conducting a literature survey to extract all this related information is time-consuming and a laborious process and requires significant switching between different resources. Another significant issue is the miRNA-based publications have been growing exponentially (Figure 1), making it difficult for existing miRNA resources to be up to date.

**Figure 1:**
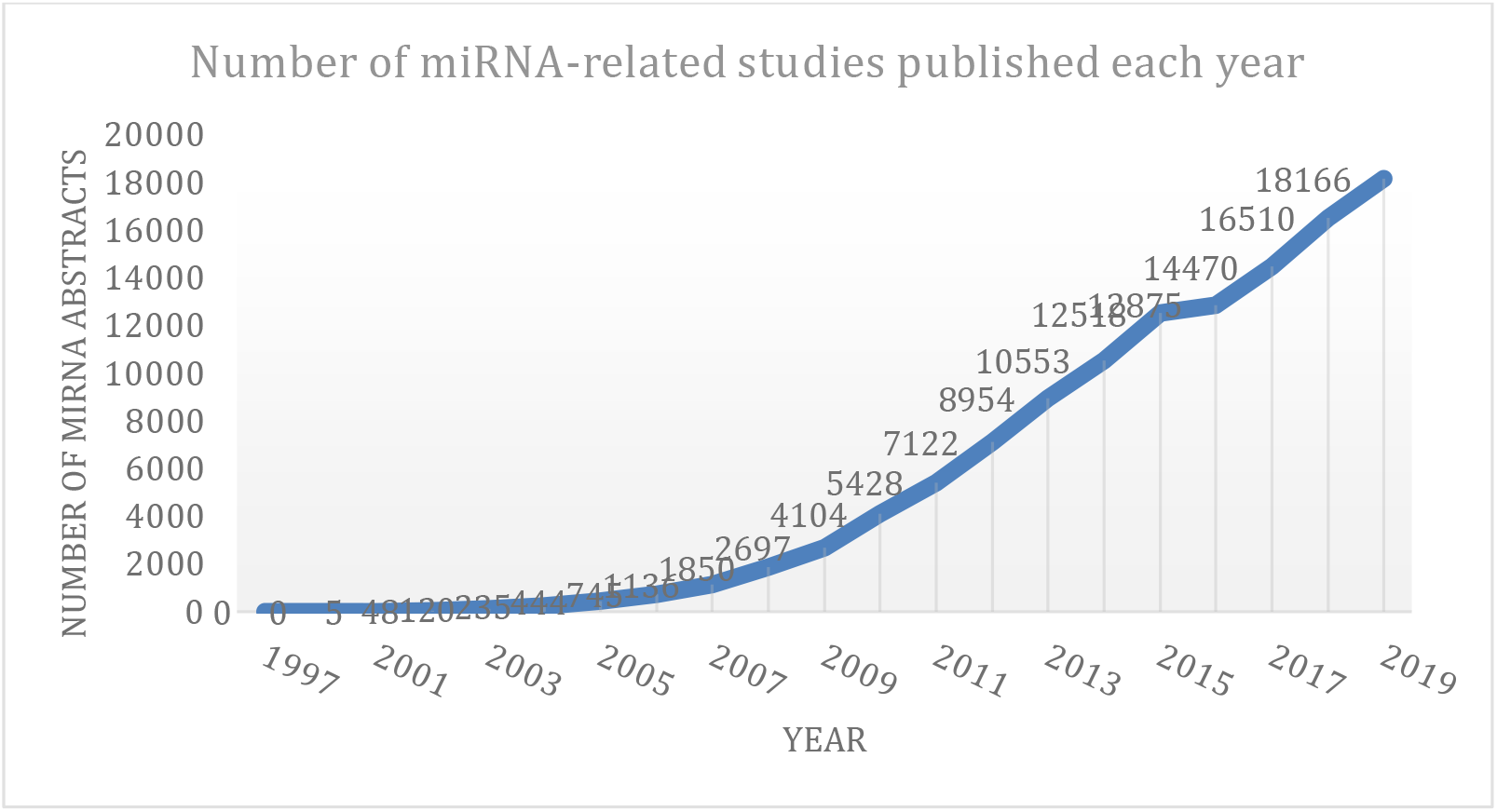
Exponential growth of miRNA publications obtained from Medline using keyword “miRNA” OR “microRNA”.

As such, there is a critical need for resources that can significantly reduce the cumbersome information retrieval process and quickly obtain relevant and integrated information from miRNA-related studies. For this reason, we designed emiRIT (extracting miRNA Information from Text), which mines different miRNA information from PubMed abstracts published between January 1997 to May 2020 (11). Our resource combines the mined results of several text mining tools on a large scale in one place with a unified output format. Thus, our resource offers a variety of miRNA information in one location, making the navigation between different information easier. To include the most widely studied miRNA aspects, we focus on recognizing biological entities (bioentities) of type i) *miRNA*, ii) *gene*, iii) *disease (currently only cancer),* iv) *biological processes and pathways,* and v) *extracellular locations* (transporters and biofluids). Bioentities are linked to publicly available standard ontologies/databases to ensure smooth querying, sorting and filtering capabilities in our interface and expand querying abilities to integrate with external resources. The text mining tools are applied simultaneously on miRNA literature to provide consistent and regular updates from new papers. Since all the information are mined from abstracts, we are able to link all our extracted results to the literature. Currently, many of the existing resources do not have direct link to the literature evidence, hindering a full interpretation of the miRNA information due to lack of context.

A unique aspect of emiRIT is that it provides a more detailed picture of a miRNA’s role in the context of a disease. The different roles of miRNAs we detect in diseases are illustrated by the following examples:

1. Role in disease process and outcome: *“High miR-21 expression is associated with poor survival and poor therapeutic outcome.”* [PMID: 18230780 (12)]
2. Role in disease treatment: *“The role of miR-181a in conferring cellular resistance to radiation treatment was validated both in cell culture models and in mouse tumor xenograft models.”* [PMID: 22847611 (13)]
3. Role as biomarker: *“Low-level expression of microRNAs let-7d and miR-205 are prognostic markers of head and neck squamous cell carcinoma”* [PMID: 19179615 (14)]
4. Role as therapeutic target: *“These findings suggest that miR-24 could be an effective drug target for treatment of hormone-insensitive prostate cancer or other types of cancers.”* [PMID: 20195546 (15)]
5. Unspecified role in disease: *“Altered expression of miR-21, miR-31, miR-143 and miR-145 is related to clinicopathologic features of colorectal cancer.”* [PMID: 18196926 (16)]
6. Differential expression in disease: *“Extrapolation of this study to human primary HCCs revealed that miR-122 expression was significantly (P = 0.013) reduced in 10 out of 20 tumors compared to the pair-matched control tissues.”* [PMID: 16924677 (17)]

The mined information is stored in a database and presented to users through an interface. On emiRIT’s interface, users can examine different miRNA aspects in one place, smoothly navigate between various aspects for a broader understanding of miRNAs’ role, and also narrow down the information to a specific biological context. Our main goal is to provide an up to date and user-friendly resource to facilitate access to relevant miRNA information from the literature. In the remainder of this paper, we discuss related work, followed by the description of the pipeline to extract, store and present miRNA information comprehensively mined from Medline abstracts.

## 2. Related Work

As discussed in the previous section, miRNA data are scattered across multiple publications and databases. Several of the databases shown in Table 1 are literature-based and curated by experts, which makes these resources high quality but hard to maintain and keep up with the most recent results. The last update of several of databases in the table dates back to more than 5 years ago.

**Table 1:** A sample of existing literature-based miRNA databases

| Category | Resource Name | Short Description | Year of last update |
| --- | --- | --- | --- |
| miRNA-target | DIANA-TarBase (18) | Validated miRNA-target interactions | 2018 |
|  | miRWalk (19) |  | 2020 |
|  | miRTarBase (20) |  | 2018 |
|  | miRecords (21) |  | 2013 |
| miRNA-Transcription Factor | TransMir (22) | Validated Transcription Factor - miRNA regulations | 2018 |
| miRNA-disease | miR2Disease (23) | Validated dysregulated miRNAs in human disease | 2009 |
|  | OncomirDB (24) | Validated or potentially pathogenic roles of dysregulated miRNAs in cancer | 2014 |
|  | HMDD (25) | Experimentally supported human miRNA-disease associations | 2019 |
| miRNA-pathway | miRwayDB (26) | Validated miRNA-pathway associations | 2020 |
| miRNA-extracellular locations | miRandola (27) | Validated extracellular circulating non-coding RNAs | 2017 |
| miRNA-expression | dbDEMC (28) | Differentially expressed miRNAs in human cancers | 2017 |
| miRNA-expression | PhenoMir(29) | Manually curated database collecting differentially regulated miRNA expression in diseases and biological processes | 2011 |
| miRNA-sequence,<br>miRNA-expression | miRBase (30) | Collects and construct information about published miRNA sequences and expression profiles | 2018 |
|  | miRNEST(31) | Combines sequence, expression from external database with predicted targets, mirtrons and miRNA gene structure | 2016 |

Due to the sheer amount of miRNA publications, as indicated in Figure 1, text mining methods have been increasingly adopted for automatic extraction of relations between a miRNA and a target/process/disease to assist in tasks such as database curation and knowledge discovery (miRNEST (31), miRSel(32), miRTex(33), miRCancer(34), miRiaD(35), Murray *et al*., 2010 (36), DES-ncRNA(37)). Except for a few, most of the approaches are limited by their narrow scope. DES-ncRNA uses only simple co-occurrences within sentences to find miRNA connections but lack the robustness to capture connections beyond co-occurrence. Work done by Murray *et al.,* 2010 is network oriented. They do not directly find a miRNA’s connection to a disease or process. Instead they combine a miRNA’s connection to genes and then use curated databases to get a gene’s connection to diseases and processes to finally generate a Cytoscape network.

emiRIT, on the other hand, focuses on extracting connections between a miRNA and a gene/disease/process/extracellular location by utilizing text mining tools that capture patterns from the syntactic structure of sentences. The following section describes the pipeline for development of emiRIT.

## 3. System Design

This section describes the design and structure of the two major components: - a database to store relevant miRNA information and an interface to interact with the stored data.

### 3.1 Database

#### 3.1.1 Database Content

To meet the desired needs of our resource, the database stores the following:

- Various **relations involving miRNA** that are extracted from text in publications.
- The **biological context**, such as a disease context or a process context, in which the above extracted relations were mentioned to provide more perspective to specific user needs.
- The **literature evidence** for these miRNA relations so that users can read the sentence or the whole abstract to interpret the information and get a complete understanding of the relations in a fuller context.
- **Standardized/normalized text mentions** of all entities allowing for extended querying capabilities and smoother integration with external resources and ontologies leading to a more connected resource. Standardizing and normalizing entities using pertinent ontologies will also provide access to additional descriptions for these entities.

#### 3.1.2 Creation of the Database

In this section, we will describe the processing of text from scientific publications as well as discuss the recognition and normalization of entities and the extraction of the relations between them. Figure 2 shows the workflow of how the database is created using abstracts and viewed in the interface.

**Figure 2:**
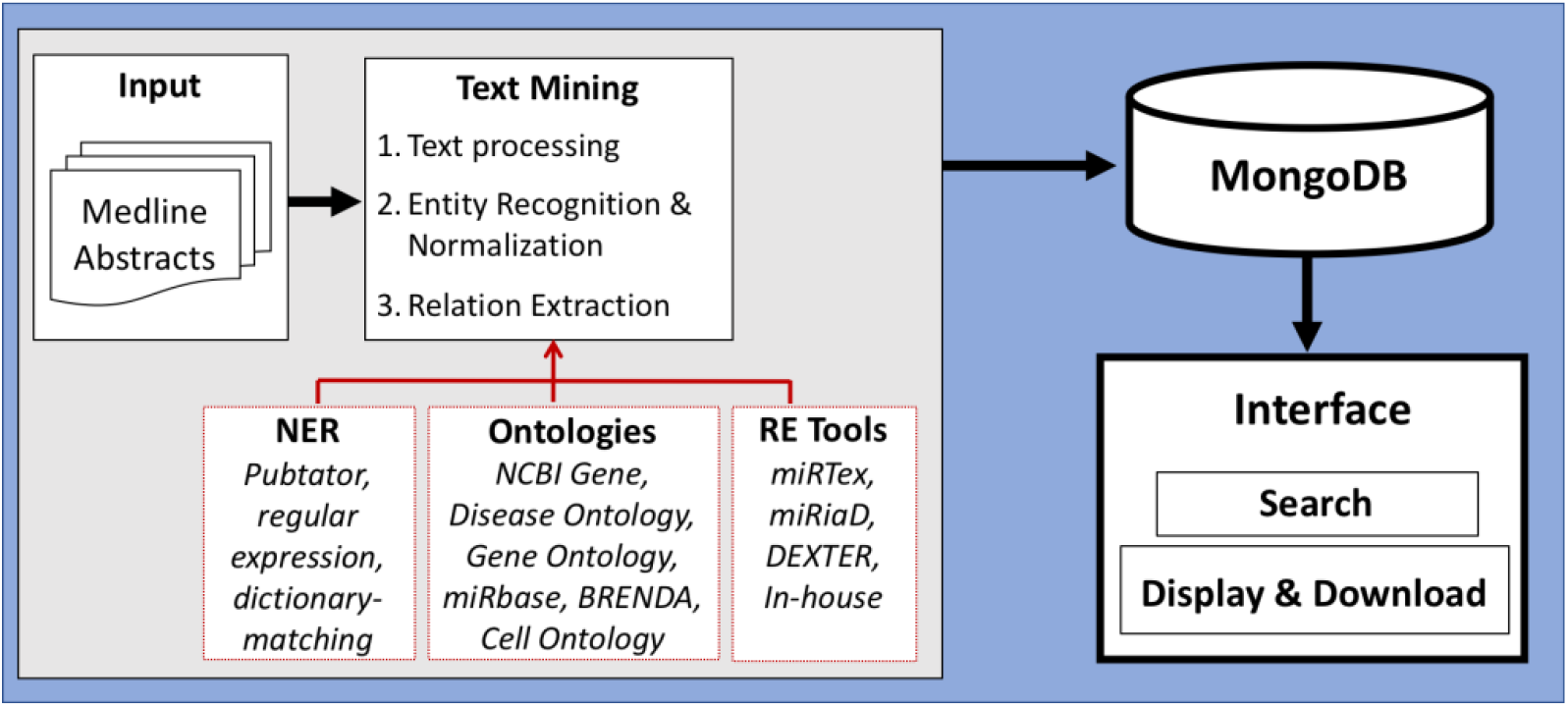
Workflow of creation of database by processing and storing miRNA-relevant information and viewing through an interface.

##### 3.1.2.1 Text Preprocessing

To support the extraction of the relations and provide literature evidence, we store miRNA-related abstracts from PubMed split into sentences in the database. We download abstracts mentioning miRNAs from Medline using the query “*miR OR miRNA OR microRNA*” on PubMed. We split each abstract into sentences using the Stanford CoreNLP sentence splitter (38).

##### 3.1.2.2 Entity Recognition and Normalization

As discussed earlier, we will focus on the following types of bioentities in our miRNA resource: miRNAs, genes, diseases, biological processes and pathways, and extracellular locations (transporters and biofluids). For miRNA detection, we use a regular-expression-based in-house tool (see Supplementary Table S1 for the regular expressions). The detected mentions are normalized to the corresponding family Id in miRBase (39). For genes and disease mentions, we use PubTator (40) to detect the mentions. PubTator normalizes genes to NCBI Gene Ids and diseases to MESH Ids. We keep the same normalization for genes, but we map these MESH IDs to Disease Ontology Id (DOID) (41) using the publicly available mapping table from http://purl.obolibrary.org/obo/doid.obo. For other entities, we develop our own dictionary-based matching technique to obtain the closest match.

For biological processes, the dictionary is created using a combination of terms and their synonyms from Gene Ontology (GO) (42), and terms mined from text using patterns. For example, on encountering text such as “cellular processes such as migration, invasion, and cell death”, we extract the three listed process terms. Commonly appearing terms mined from large amounts of Medline abstracts were thus collected. The process terms are normalized to GO accession numbers, whenever matched. Pathway mentions are detected and normalized to a list of pathways from the Pathway Ontology. For extracellular locations, we use exRNA forms and fluid samples from miRandola (27) to build our dictionary and normalize the fluid mentions from the text using Brenda Tissue Ontology (43).

##### 3.1.2.3 Relation Extraction

This subsection discusses the extraction of various relations stored in the database

###### miRNA-Gene

One of the main relations we capture are between a miRNA and a gene since miRNAs are important gene regulators. To capture such a relation, we use a text mining tool called miRTex (33), which detects the miRNA and its target gene (a gene regulation of a miRNA). miRTex also detects where a miRNA regulates gene expression - either indirectly or when it is not clear if the regulation is a direct result of targeting. For example, from the following sentence alone, we cannot infer if the regulation is due to a single step:

###### Example 1

“*Mechanistic studies disclosed that, <u>miR-340</u> over-expression suppressed several oncogenes including <u>p-AKT</u>, <u>EZH2</u>, <u>EGFR</u>, <u>BMI1</u> and <u>XIAP</u>”* [PMID: 25831237 (44)]

miRTex has been evaluated and found to be a robust extraction system of the three different types of miRNA-gene relations from abstracts and full-length articles with high precision, recall and F-scores close to 0.9.

###### miRNA-Process

The next relation we focus on are between a miRNA and a process since most biological processes are subjected to miRNA-dependent regulation. For this purpose, we have extended another text mining tool, miRiaD (35). Central to miRiaD is the detection of the “involvement”, “regulation” and “association” between a miRNA and a disease aspect For our resource, we extend the Connections with Association, Involvement and Regulation (CAIR) framework of miRiaD to make the connections between miRNAs and processes and pathways, irrespective of the presence of disease terms in the abstract. In the example below, the CAIR framework will detect that miR-29b positively regulates apoptotic process.

###### Example2

*<u>MicroRNA-29b</u> promotes high-fat diet-stimulated endothelial permeability and <u>apoptosis</u> in apoE knock-out mice by down-regulating MT1 expression*. [PMID: 25131924 (45)]

###### miRNA-Disease

As mentioned before, the varied roles of miRNA in diseases have been widely researched, including their role as potential biomarkers and therapeutic targets, their impact on the treatment of diseases and disease outcomes. Instead of just stating that there is an association between a miRNA and a disease, we seek to present a more detailed picture of the role of a miRNA in context of disease by distinguishing between these different roles. Specifically, these roles are i) impact of miRNA on disease process and outcome, ii) influence on disease treatment iii) diagnostic role as biomarkers, iv) role as therapeutic targets in diseases, v) others, where the particular role is not clear, but the miRNA is associated with a disease or regulates a disease.

Since miRiaD was developed to capture the different ways a miRNA is linked to a disease or a disease aspect, we extend miRiaD significantly for our purpose. For the first two relation types, we use the CAIR framework from miRiaD whereas for the next two relations, we use a group of rules clubbed together to form the “is_A” framework that can capture relations such as “X is a Y”, or “X acts as Y”, or “X serves as Y”. While miRiaD clubbed all the five different roles together and called them “disease aspects”, we have enhanced miRiaD’s ability to take the arguments of the relations and we separate the disease aspects based on the type of the arguments. As an illustration of argument-based separation, examples 3 and 4 below show how a sentence depicting a miRNA’s role as biomarker and therapeutic target are structured.

###### Example 3

*“From a clinical point of view, our study emphasizes <u>miR-122</u> as a <u>diagnostic and prognostic marker</u> for HCC progression*.” [PMID: 19617899 (46)]

###### Example 4

“*Our data suggest that <u>miR-429 may serve as a potential anticancer target for the treatment</u> of HCC”*. [PMID: 28440423 (47)]

In example 3, the “is_A” framework captures “miR-122” is_A “diagnostic and prognostic marker” while the same rule captures “miR-429” is_A “anticancer target for the treatment” in example 4. In both cases, we look at the type of the arguments of the relation and separate them into “miRNA is a biomarker” and “miRNA is a therapeutic target”. miRiaD had reported a high recall and precision with an F-score close to 0.90 when evaluated on a curation task as well as for general extraction of miRNA to disease associations.

Finally, knowing which miRNAs are differentially expressed in disease is important, especially for understanding or generating hypotheses about the underlying causes. To capture the up or down-regulation expression of miRNAs in disease vs non-disease states from research literature, we use a tool called DEXTER (48). DEXTER was designed after an extensive study of textual mentions of comparisons (49). It detects the differential expression levels as well as the location of the expression levels such as in cell lines or tissue samples, patient groups, control and others. DEXTER was evaluated and precision greater than 0.90 with an F-score close to 0.80 was reported for general extraction of differentially expressed genes and miRNAs in the context of diseases. In the Section discussing view of miRNA-aspects in the Interface, Figure 8 shows how the interface will display the different miRNA roles extracted using miRiaD and DEXTER to cater to the information needs of our resource. Since both miRiaD and DEXTER were developed specifically for cancer, we have currently restricted emiRIT to only cancer with plans to extend to other diseases in future.

###### miRNA-extracellular locations

miRNAs are increasingly being studied as potential biomarkers of diseases because of their abundance and stability in extracellular fluids, transported via membrane-bound vesicles such as exosomes, or complexed with high density lipoprotein (50, 51). Hence, we focus on extracting information about miRNAs in extracellular locations. We use the Extended dependency graph (EDG) (52) framework to capture the syntactic structure of sentences and extract direct relations between a miRNA and biofluids, such as tear, serum, plasma and others, or extracellular transporter forms, such as vesicles, exosomes, protein complexes, and others. As a first step, we use the EDG framework to capture simple patterns that are focused on high precision, for cases where a miRNA and an extracellular location appears in close textual proximity. For the second step, we focus on capturing cases where the miRNA and the extracellular location are not in close textual proximity, but the the extracellular location is explicitly mentioned in experimental context of the paper. Examples 5 and 6 show the two different types of cases we focus on.

###### Example 5

*“After qRT-PCR validation, only one <u>seminal plasma</u> miRNA, <u>let-7b-5p</u>, was found significantly decreased in severe asthenozoospermia cases compared with healthy controls.”* [PMID: 29653228 (53)]

###### Example 6

[PMID: 32373058 (54)]

a. *We analyzed the expression of three microRNAs in <u>serum</u> of 18 patients (DMD 13, BMD 5) and 13 controls using droplet digital PCR.* [Sentence 2]
b. *We found that levels of <u>miR-30c</u> and <u>miR-206</u> remained significantly elevated in DMD patients relative to controls over the entire study length*. [Sentence 7]

For the second step, we implement the Patient Context (PC) sentence detection from eGARD (55), which provide information about the patients involved in the study. Our assumption, following the study of at least 20 abstracts, is based on the fact that any extracellular fluid samples from these patients are highly likely to have a connection to the miRNAs being explored in the same paper. Using this new relation extraction tool, we were able to extract 3782 miRNA-extracellular location pairs using the EDG framework and 2173 pairs using the PC sentences. We sampled about 100 abstracts from our database that contained mentions of miRNA and extracellular locations. From these 100 abstracts we were able to find 136 miRNA-extracellular location pairs using both the EDG framework and PC sentences. We manually checked each of these 136 pairs and found 133 of them were indeed correctly paired whereas 3 of them were paired incorrectly. We plan to improve our new miRNA-extracellular location relation extraction tool in future and conduct further evaluation by comparing the results from the tool to a manually annotated dataset.

#### 3.1.3 Database Structure

Based on our previous experience of providing access to information stored in a database to users via an interface in iTextMine (56), which is an integrative text mining system for knowledge extraction developed in our lab, we choose to store our data using a standardized JSON format (57). This format is a lightweight data-interchange text-based format commonly used for transmitting data in web applications. Our data is centered around miRNA-relevant abstracts that undergo various text processing to retrieve the entities and their relations within each abstract. This document-centric data is then stored in a non-relational database. We use MongoDB (58) since it can accommodate diverse types of data, including documents, and the abstracts can be easily represented using a JSON format and then directly inserted into the database as a document collection. Figure 3 shows an example of how an abstract is represented in the database. An example of the JSON format of a document is provided in Supplementary Figure S2. The pipeline for creating the database also includes an “Update” step, which will ensure that the database is brought up to date every few months and the most current miRNA information is captured in our resource.

**Figure 3:**
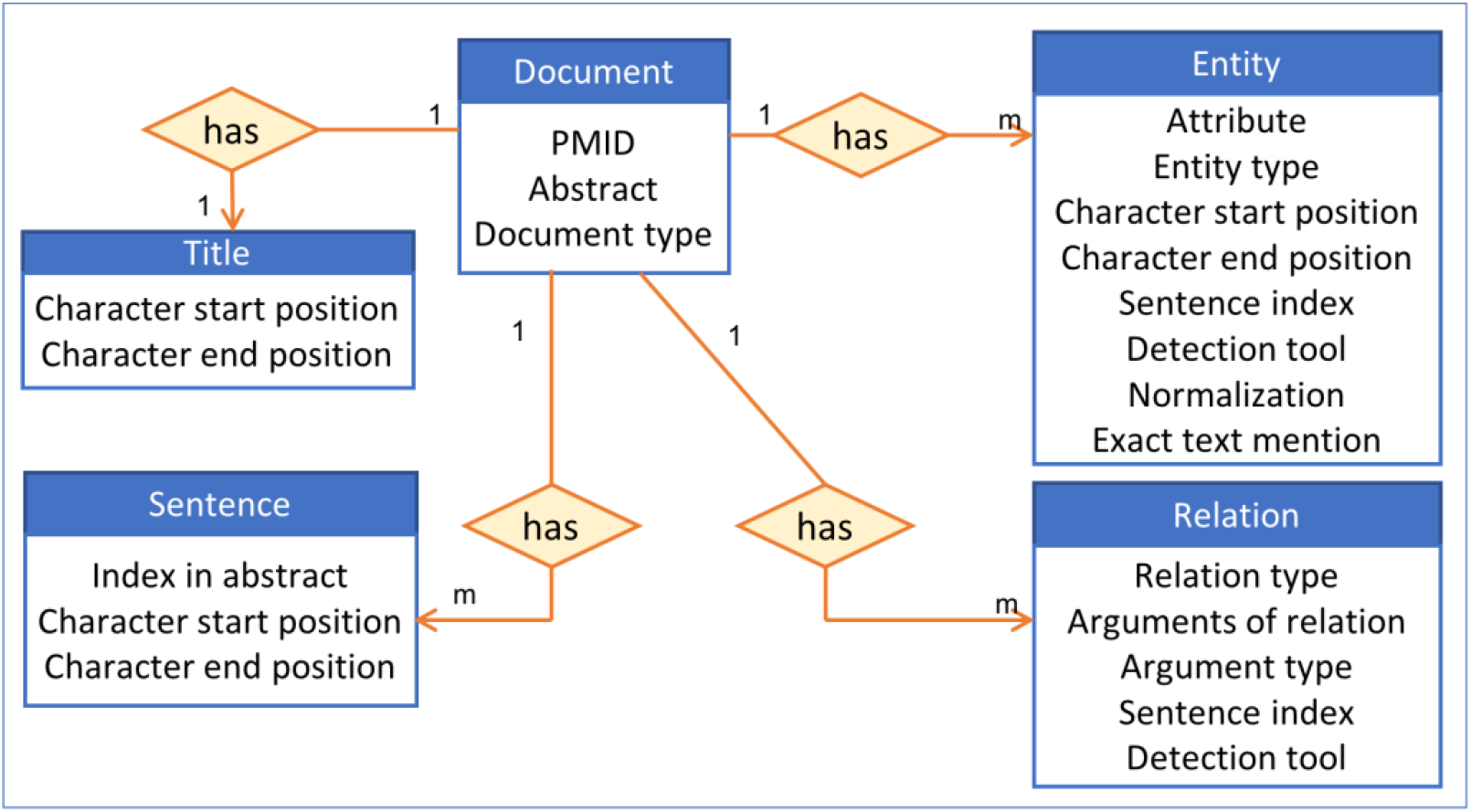
High-level view of the information stored in the database for an abstract

### 3.2 emiRIT Interface

We created the interface https://research.bioinformatics.udel.edu/emirit/ using Flask 1.0.3 and Bootstrap 4.0. The various functionalities of the interface are described in the following subsections.

#### 3.2.1 Search Mode

There are two main modes of querying the database through this interface. The first type is the miRNA-centric search, where users can observe the different connections between a specific miRNA stored in the database and other entities. The second type is a context-centric search, where users can observe the different miRNA connections in a certain context using a general keyword query. Unlike, the miRNA-centric search, the context-centric search is similar to a PubMed search. Here, we can include separators like AND/OR in our query. To every query, we explicitly add “miR” OR “miRNA” OR “microRNA” and use the resultant query to search in the NCBI PubMed database and retrieve a list of PMIDs. We then search our database using this list of PMIDs and extract the various information for the common PMIDs from our database.

Since most of the miRNA research are conducted in the context of a disease, we provide a specialized context-centric search, which limits the context to a specific disease. Users can search the database using the DOID or official DOID name and observe the different connections between miRNA and other entities in the context of the corresponding disease. The DOID is used to retrieve the disease name and its synonyms and using a combination of these disease terms and miRNA terms, a query, similar to the general context-centric search, is constructed to search in the NCBI PubMed database. Currently, we narrow our query to only Cancer-specific diseases. Figure 4 below shows a screenshot of the different search modes in our interface.

**Figure 4:**
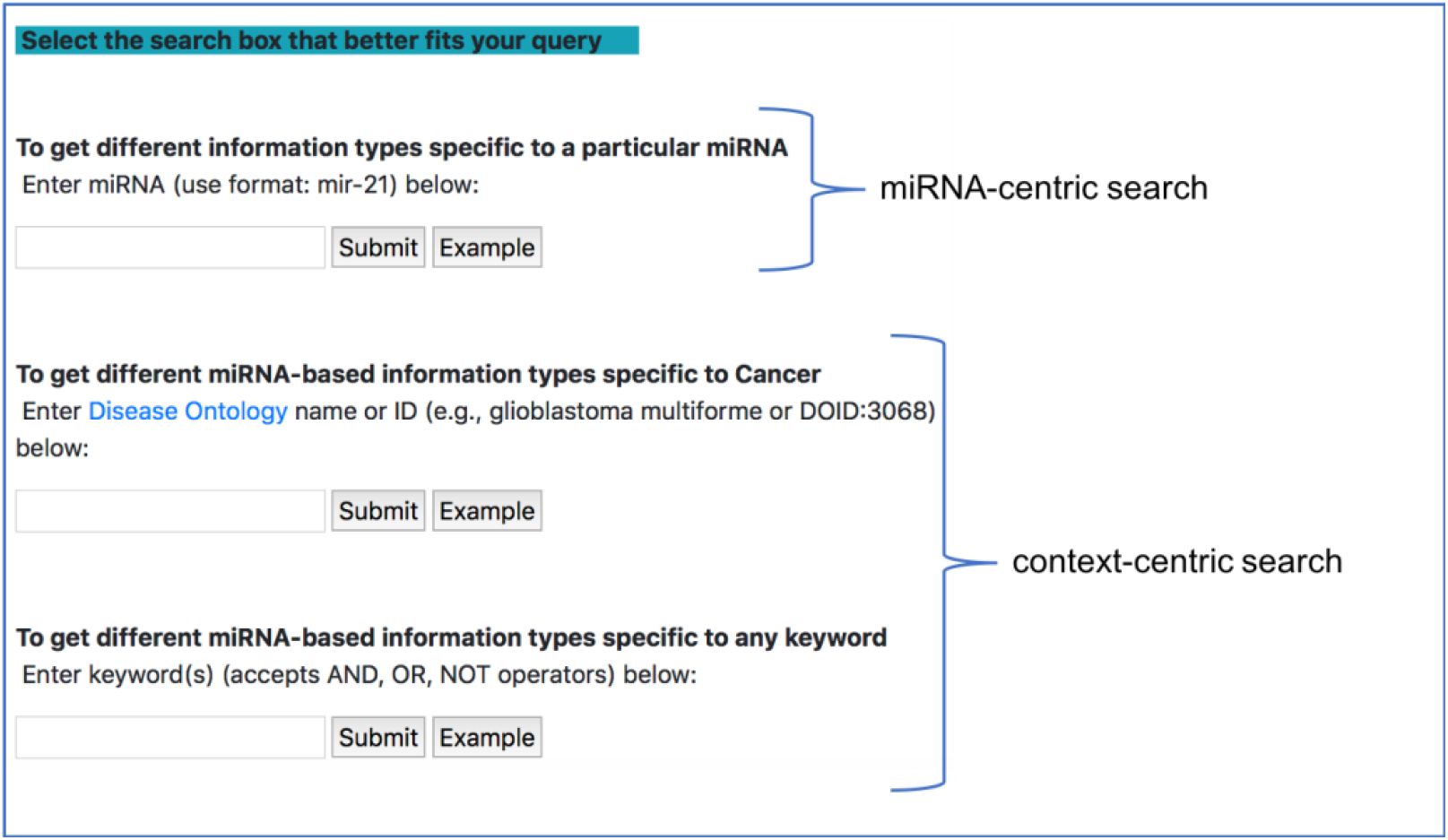
miRNA-centric search and context-centric search mode in the interface

#### 3.2.2. View of miRNA-aspects

##### High-level view

The resulting page for any of the search queries shows how many documents were retrieved and how many entities were found. A table summarizing the different entities in each document is displayed at the bottom of the page (as shown in Figure 5). This table shows a high-level view of genes, diseases, processes and pathways involved in a relation with miRNAs for each document. These entities were found to be in a relation with miRNAs by the relation extraction tools we have discussed before. From the search result page shown in Figure 5, users can either navigate to specific aspects of miRNAs or to a specific document, as described in the following subsections.

**Figure 5:**
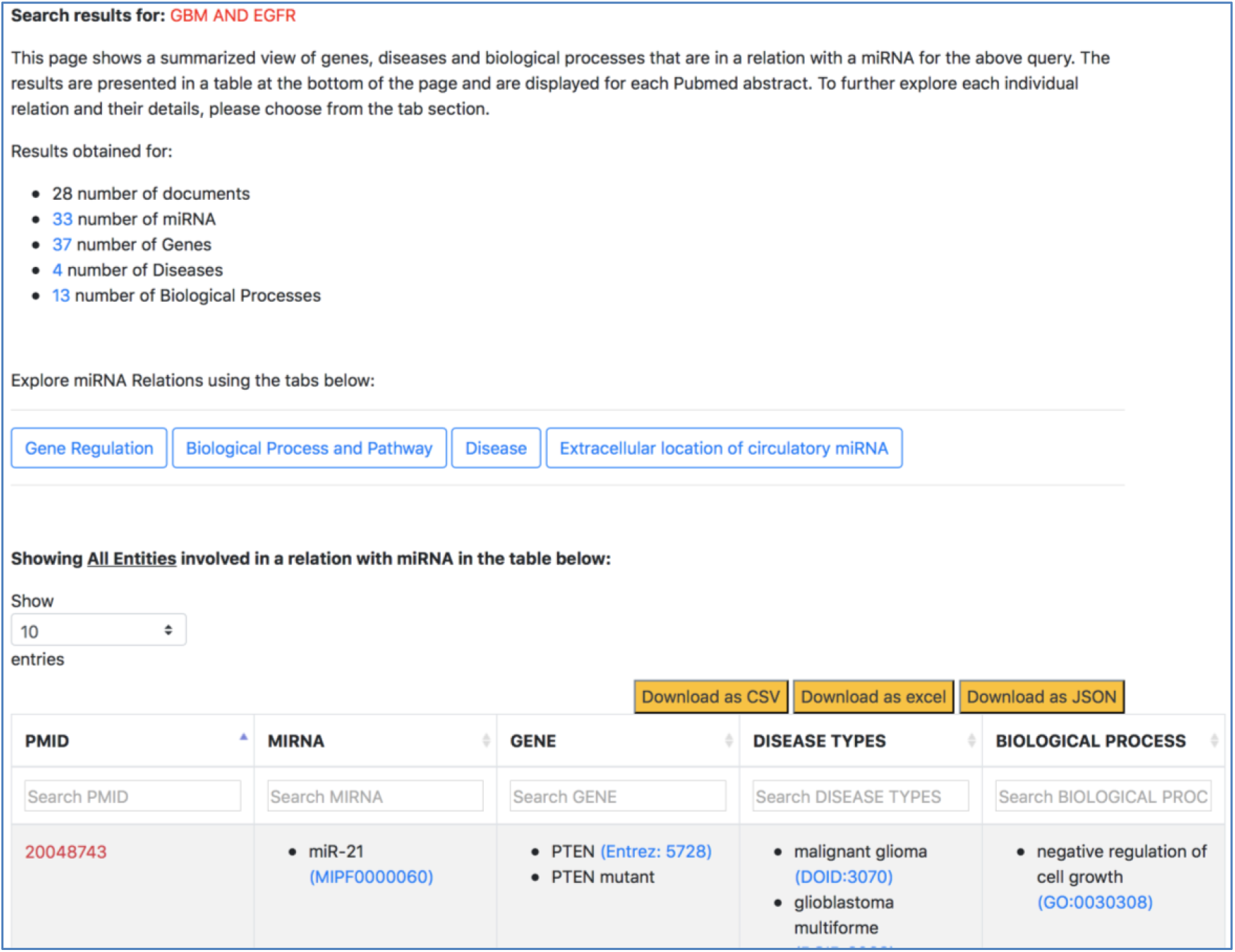
Screenshot of response page for context-centric query “GBM AND EGFR”

##### Aspect-specific view

The aspect-oriented information can be viewed by exploring the tabs at the top of the search result page (refer to Figure 6). For instance, if the user wants to know what different genes are targeted by the different miRNAs in the context of GBM, they can choose the specific “Gene Regulation” tab (Figure 6). The resulting page (refer to Figure 7) will show the targets of a miRNA in the context of GBM along with the PMID of the abstract as literature evidence. The “Disease” tab (Figure 6) will take us to another page that separates the different disease-oriented information, as shown in Figure 8. Users can look at which miRNAs are up-regulated or down-regulated in disease, what is a miRNA’s impact on the outcome of a disease or a disease process, which miRNAs were found to be potential biomarkers, and others.

**Figure 6:**
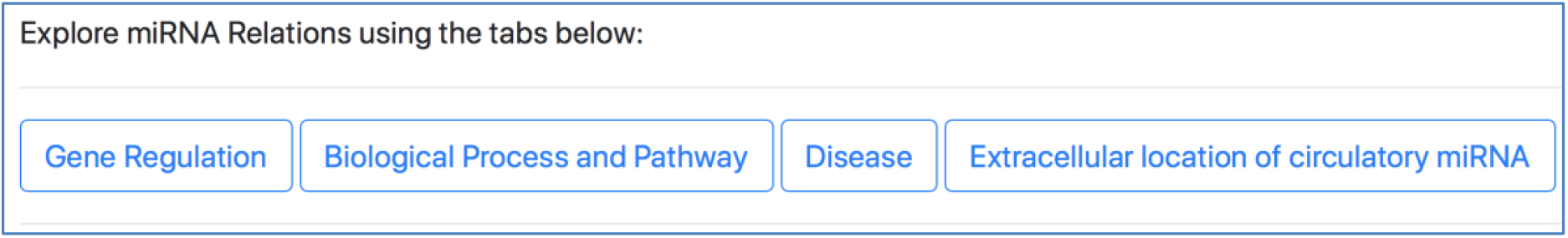
Specific miRNA relation tabs in the response page for context-centric query “GBM AND EGFR”

**Figure 7:**
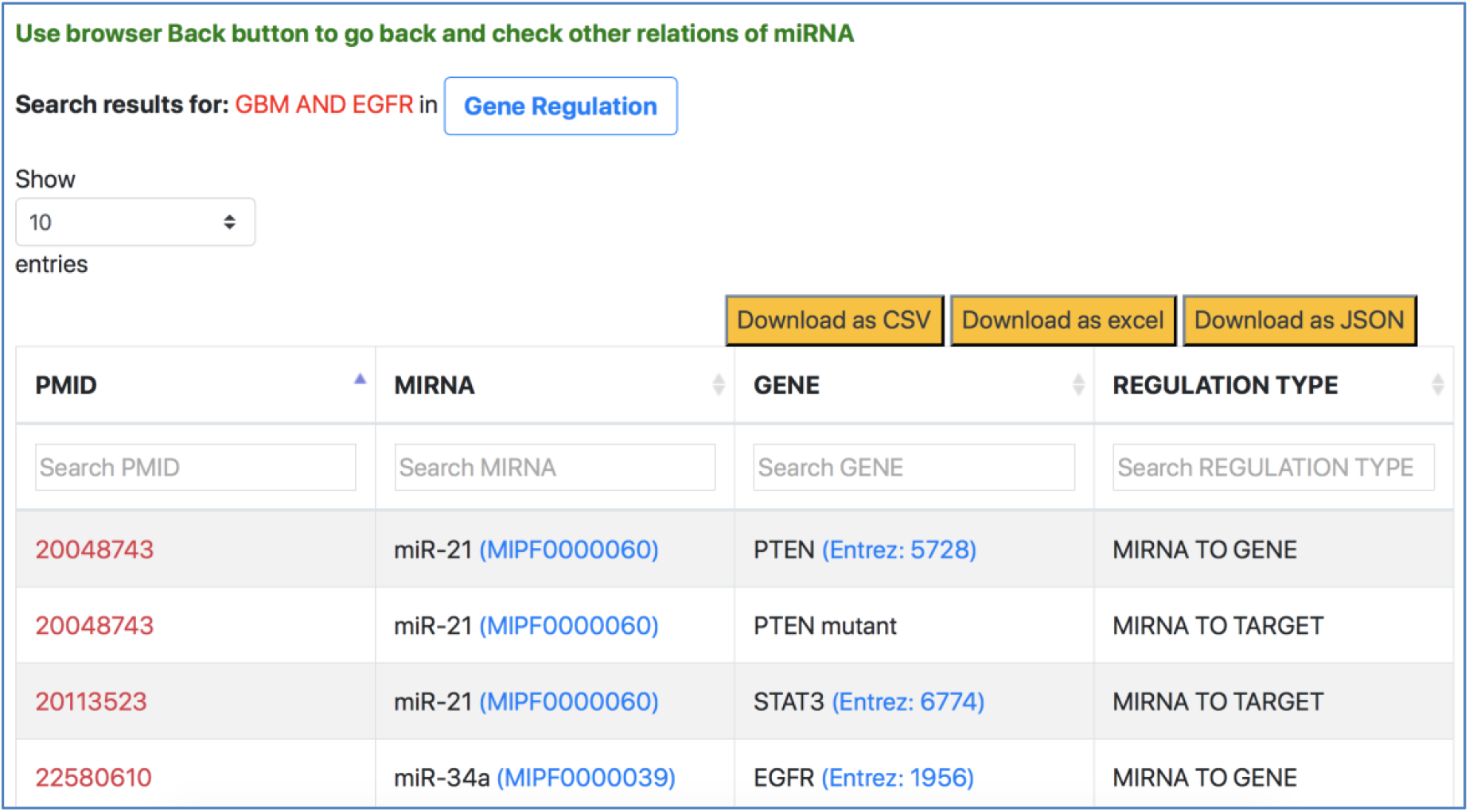
Gene Regulation information of miRNAs for context-centric query “GBM AND EGFR”

**Figure 8:**
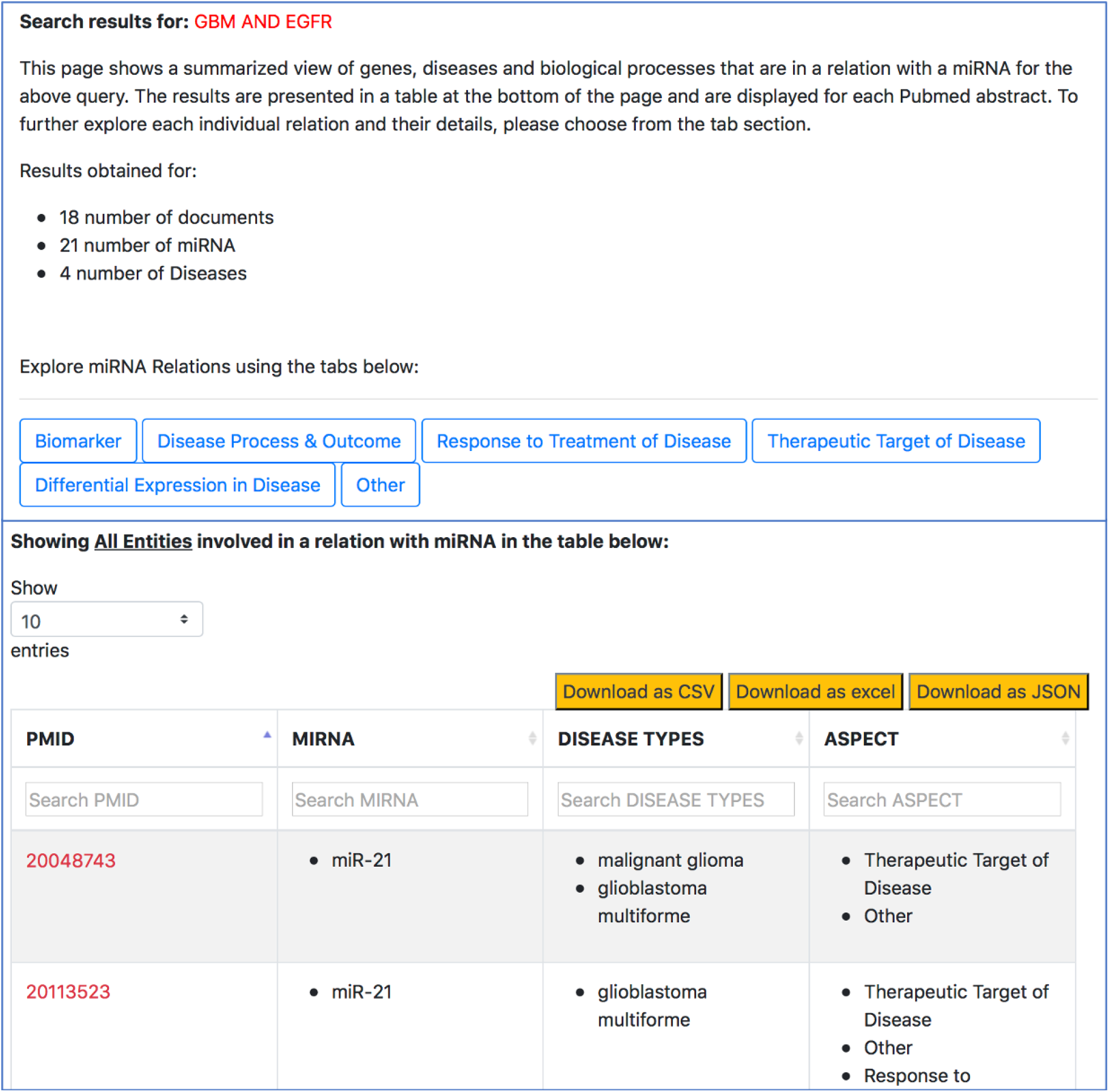
Disease-oriented information context-centric query “GBM AND EGFR”

##### Document-specific view

On clicking the PMID from any of the above pages, users will be taken to a page showing all relations about a single document. Currently, our resource only looks at abstracts since the tools we use are limited to abstracts, but we plan to extend the resource to PMC open access papers in future. As shown in Figure 9, this page shows the abstract of the document, where each sentence is separated and visible. All miRNA relations, extracted using relation extraction tools, are displayed at the bottom of the abstract. Each relation is also accompanied by the sentence number from which the relation was extracted.

**Figure 9:**
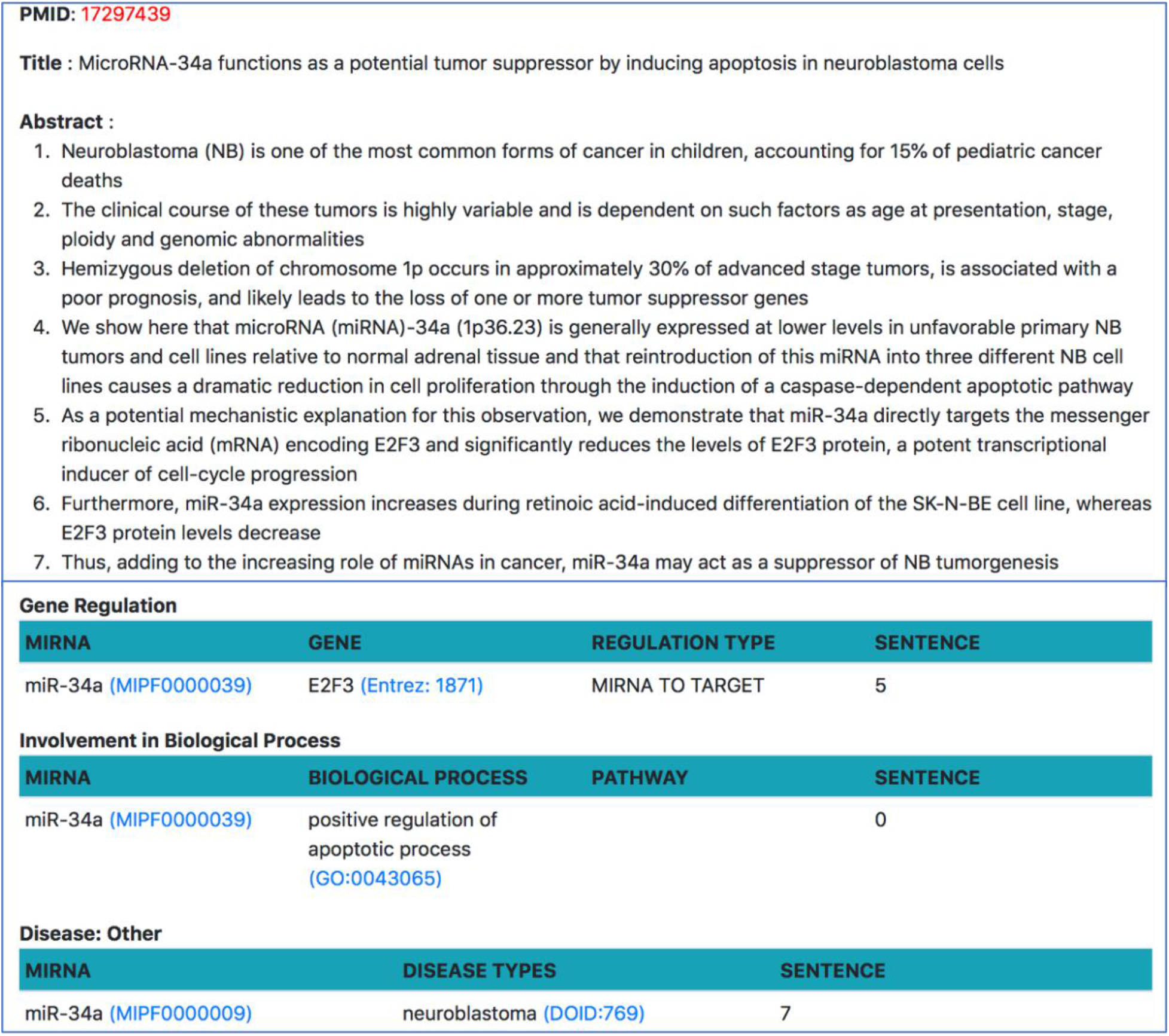
Document-specific view for PMID 17297439

#### 3.2.3 Additional features

##### Sorting and filtering capabilities

The tables in the high-level view and aspect-specific view can be sorted and filtered based on the user’s information requirement (refer to Figure 10 in Section for Case Study 1 in Results and Discussion). Sorting on the column is performed by clicking on the arrow next to the column header while filtering is performed by using the search box below the column header. The case study 1 in Results and Discussion section shows the usefulness of sorting and filtering the tables.

**Figure 10:**
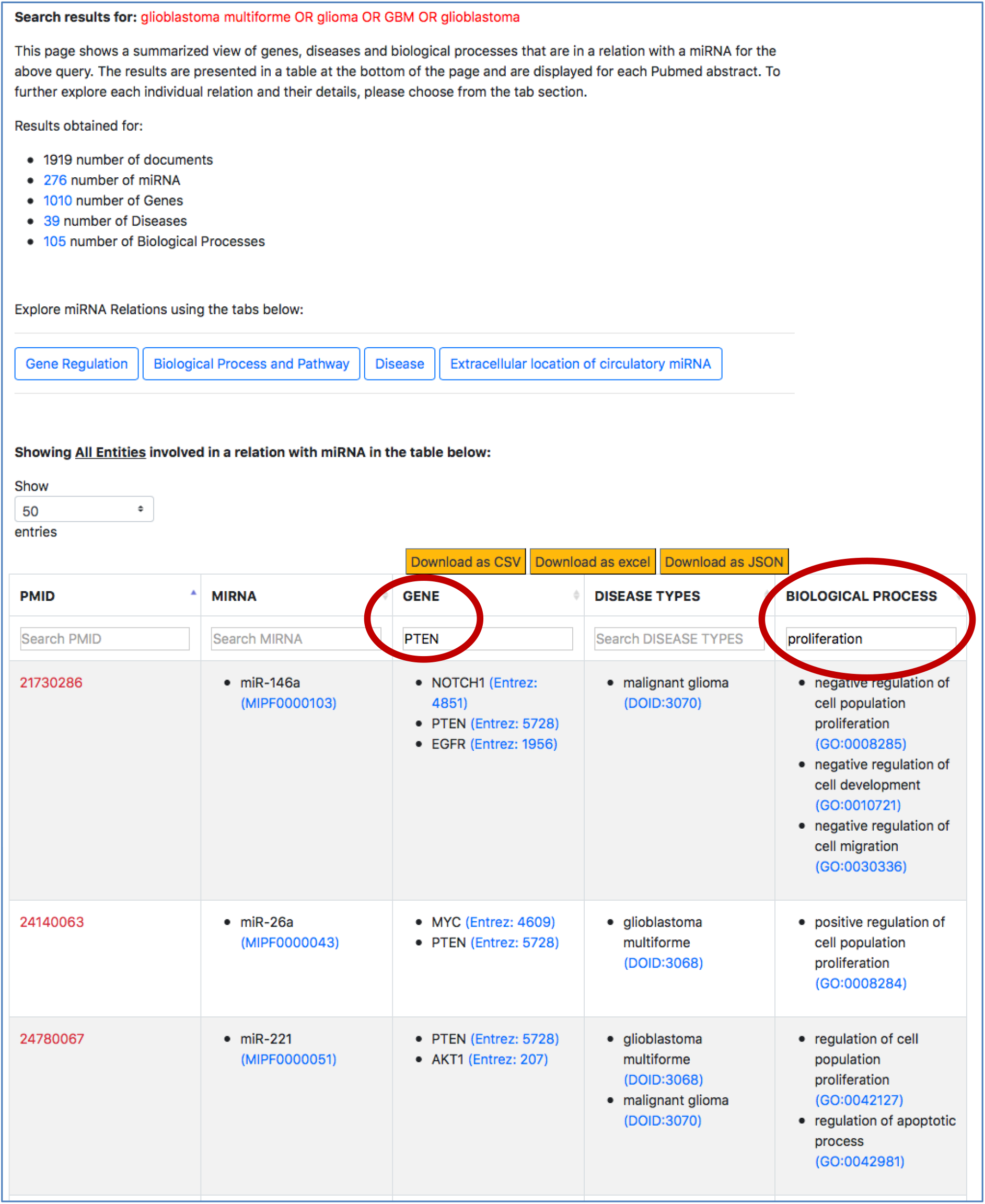
Filtered search result for “PTEN” in the context of cell proliferation in GBM

##### Ontology-driven search and link-out capabilities

We normalize the entities of different types using publicly available and standard ontologies/databases, specifically to i) ensure querying, sorting and filtering capabilities in our interface do not miss synonym of terms and ii) expand the information scope of this resource by integrating with external resources that provide descriptions each entity, such as genomic context of genes or sequences of miRNAs, and expand querying capabilities in other manually curated resources to broaden the understanding of the user about the role of a miRNA. For example, in Figure 7, when a user looks at the specific miRNA-gene relation and they want to know more about the miRNA’s sequence or the description of the gene, they can simply click on the entity term. A separate page on the browser takes them to an external ontology that describes the specific entities. Plus, normalizing the entities improves the filtered results from the tables. In Figure 10, the normalized gene terms ensure that different names of “PTEN”, such as “phosphatase and tensin homolog” are also captured when the table is filtered.

##### Download functionality

Users can download the results from the summarized table from the high-level view and relation-specific tables from the aspect-centric view as a JSON file, CSV file or an Excel file.

## 4. Results and Discussion

We collected around 121,371 miRNA-related abstracts from PubMed, out of which we extracted more than 149,233 miRNA-PMID pairs. 49,010 out of 121,371 abstracts contained relations between miRNAs and other bioentities. Table 2 shows the number of unique entities in these 49010 abstracts as well as the number of relations between miRNAs and other entities.

**Table 2:**
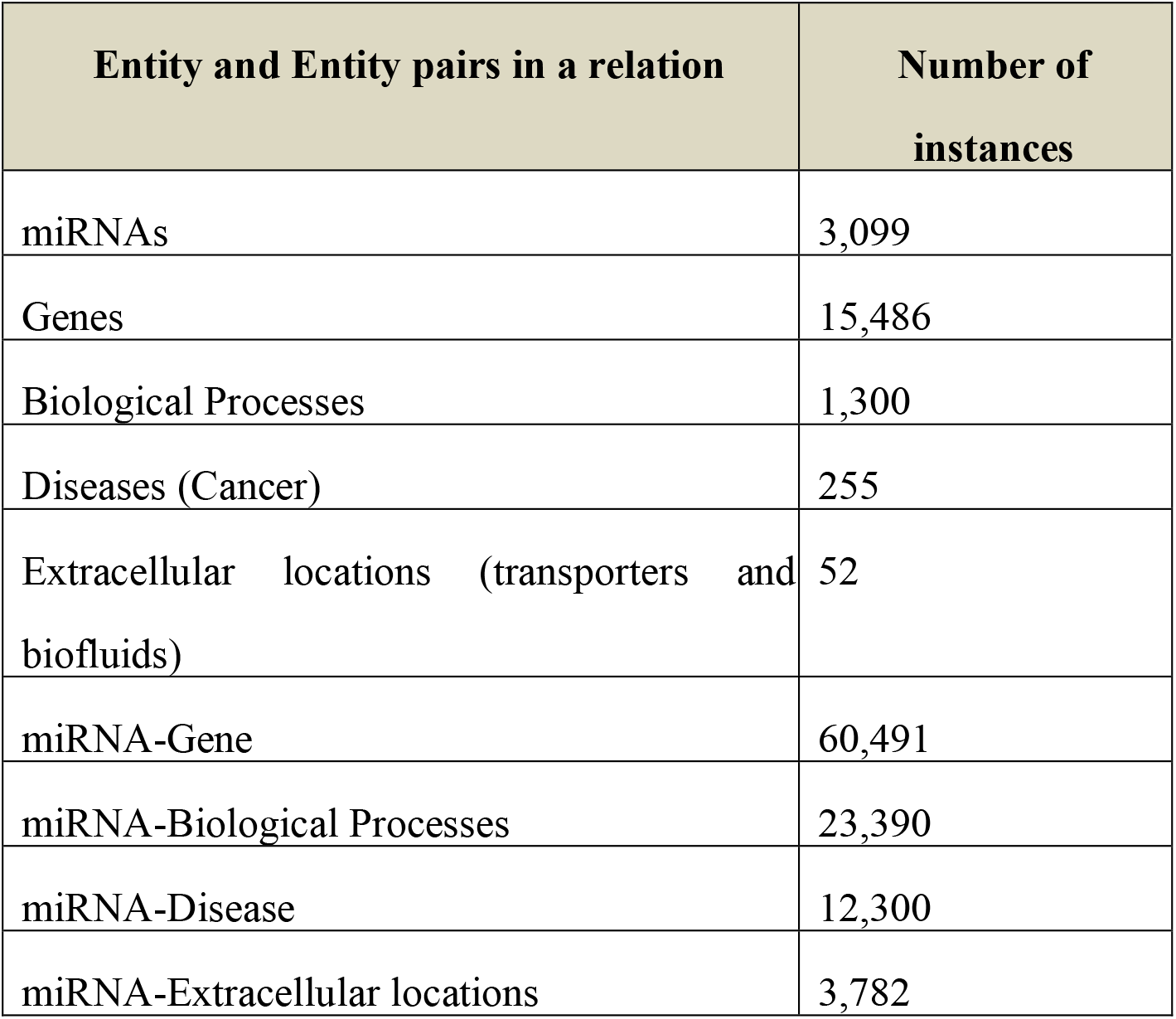
Summarized information of the number of bioentities and number of connections between miRNA and other bioentities in emiRIT.

While the number of connections express how much emiRIT extracted from the literature, it does not give us a sense of what information has been missed in abstracts. Therefore, we decided to conduct case studies to assess the information coverage of different aspects of miRNAs through our resource. Since there are no available gold standards that we can use to compare our findings, we rely on review articles for our case studies to find the extent of information emiRIT is able to capture. We assume that the users of our resource will look for miRNA information in some context. As miRNAs are mostly explored in the context of diseases, we decided to focus our case studies on two widely investigated aspects of miRNAs that are important to understand miRNA’s role in diseases – the target information and the differential expression of miRNAs. We use two highly cited review articles in these case studies, where one of them explores miRNA’s role in the context of Glioblastoma Multiforme (GBM) and the other explores differential expression in human gastric cancer. In the absence of highly cited review articles on the other miRNA aspects, we limit our case studies to the above-mentioned aspects.

### 4.1 Case Study 1: Target information of miRNAs in the context of a disease

The first case study explores the relation of miRNAs with bioentities in the context of Glioblastoma Multiforme (GBM). A recent comprehensive review explored miRNA’s role in eight hallmarks of GBM (59). We explored the first two hallmarks of GBM in our case study and compared the findings using our interface with that provided in the review.

The first hallmark question in the review investigates the aberrant miRNAs that affect Receptor Tyrosine Kinase (RTK) signaling networks to promote cell proliferation in GBM cells. The review identified 29 miRNA-target pairs described as the miRNAs regulating the targets and modulating the RTK signaling network leading to cell proliferation. We captured 25 of the 29 miRNA-target pairs mentioned in the review using our interface. We used the general keyword search with the query “(glioblastoma multiforme OR glioma OR GBM OR glioblastoma).

Following is an example of our mode of search that we conducted on the interface to retrieve 25 out of the 29 miRNA target pairs. For this example, we will consider “PTEN”, which was found to be a target for increased tumor growth by the review of oncomiRs, such as miR-17-5p, miR-19a/b, miR-21, miR-1908, miR-494-3p, miR-10a/10b, miR-23a and miR-26a. To observe how many of these miRNA-“PTEN” pairs were also retrieved using emiRIT, we filtered the aforementioned general query search result in the interface with “PTEN” as the keyword. We then observed all the miRNAs for any abstract that mentioned “proliferation”. The snapshot of a filtered table using “PTEN” in the search box above the table is shown in Figure 10.

We were able to identify miR-19a, miR-26a, miR-494-3p and miR-21 directly from the rows that contained both “PTEN” and “proliferation”. miR-23a was identified in rows that contained “PTEN” while miR-10 was found in the same row as “PTEN” and “migration”. miR-17-5p was found to target PTEN but no mention of “proliferation” or terms associated with “proliferation” were detected by our resource. Even though we were able to find miR-1908 in the context of GBM and proliferation, we could not find PTEN to be the target of miR-1908. Additionally, we were able to capture much more information than what was found in the review. In the last two rows shown in Figure 10, we not only see that miR-26a was found to be in a relation with PTEN and was involved in proliferation, we also find other miRNAs, for example, miR-221, that were associated with PTEN and proliferation. The review did not identify miR-221and PTEN as a miRNA-target pairs involved in cell proliferation. The abstract evidence, shown in Figure 11, suggests that miR-221 targets PTEN in the context of cell proliferation (sentences 6-10) in GBM.

**Figure 11:**
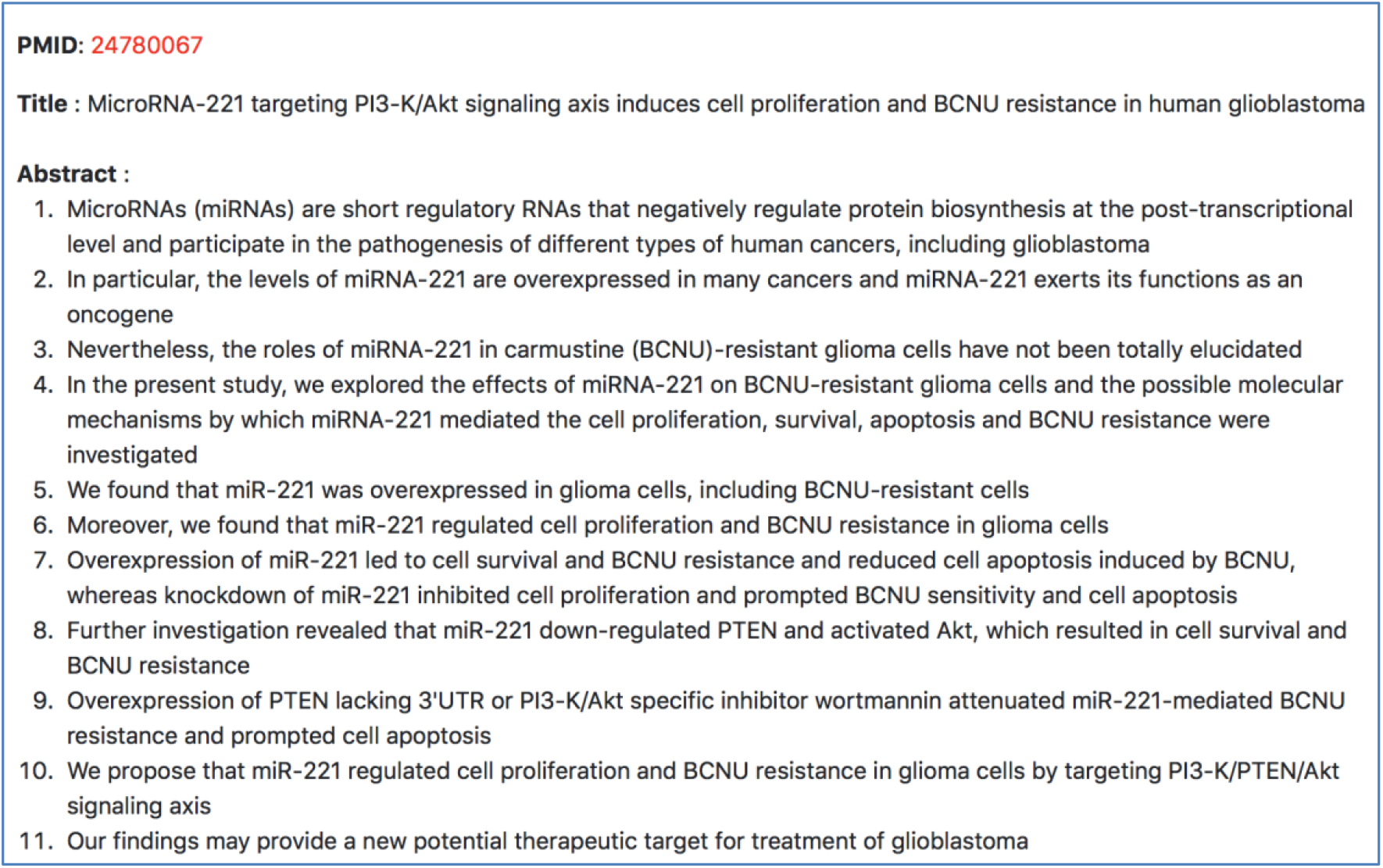
Abstract evidence for “miR-221” promoting “proliferation” by targeting “PTEN” in the context of GBM

### 4.2 Case Study 2: Differential expression of miRNAs in the context of a disease

Our second case study explores the differential miRNA expression in human gastric cancer. We used a highly cited review by Shrestha *et al.,* 2014 (60), which surveys miRNA expression profiling studies in human gastric cancer. The review mentions 41 miRNAs that were consistently upregulated and 28 miRNAs that were consistently downregulated. For our case study, we looked at these 69 miRNAs to see how many of them are found to be upregulated or downregulated using emiRIT.

On our interface, we used the disease-centric search with the query “stomach cancer” from the list of Disease Ontology diseases autocompleted by the NCBO BioPortal widget. We use “stomach cancer” because a search for “gastric cancer” in the Disease Ontology also provided results for “stomach cancer”. We then navigated to the “Differential Expression in Disease” tab from the “Disease” tab. Figure 12 shows a screenshot of a table in the resulting page from the “Differential Expression in Disease” tab. We filtered the information on this table by using the second column for miRNAs and searched for the 69 miRNAs identified in the review. Supplementary table S3 and S4 compares between our findings and that of the review for all 69 miRNAs.

**Figure 12:**
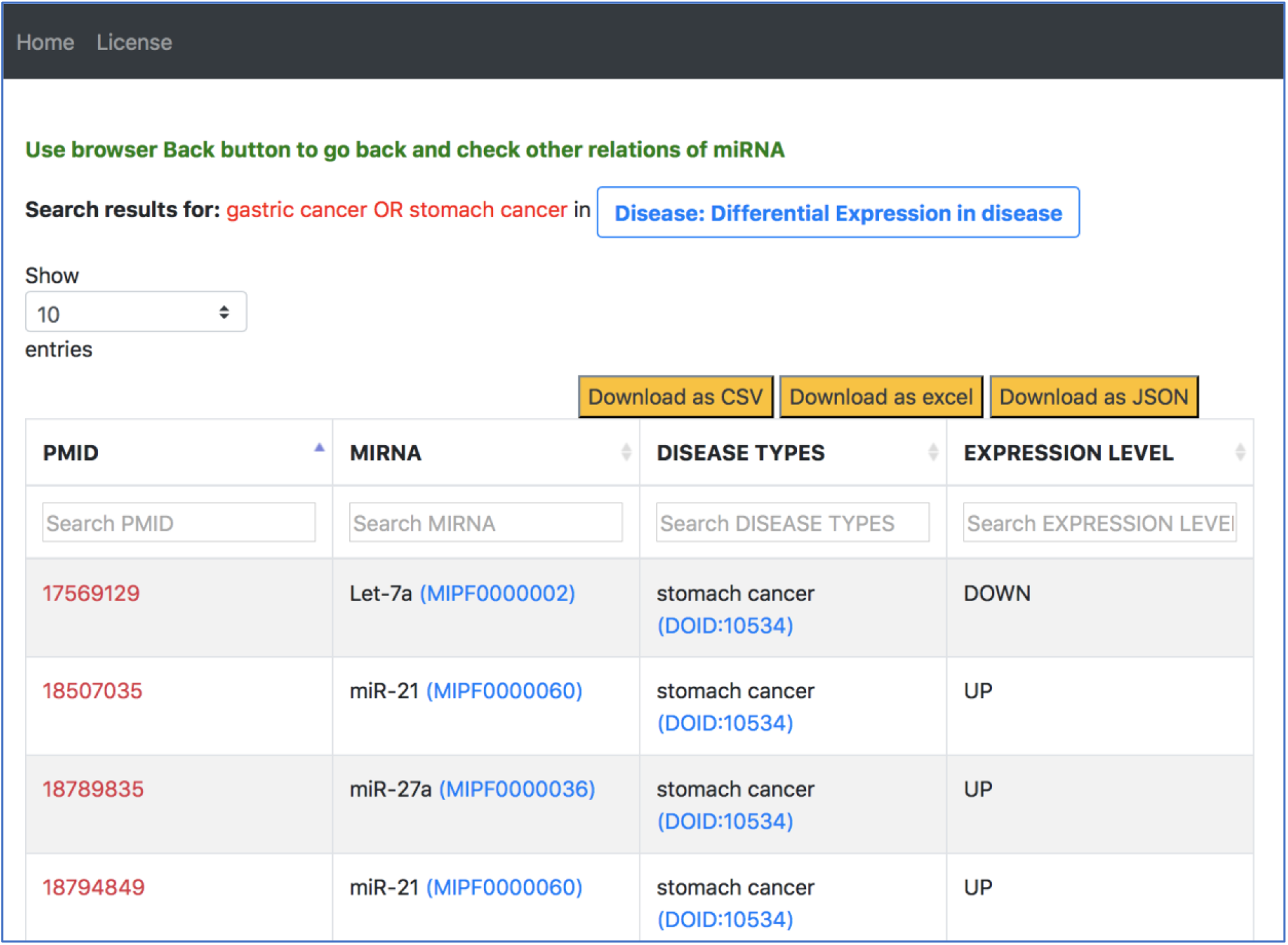
Output response page of “Differential Expression in Disease” tab for “gastric cancer OR stomach cancer” query in general keyword-centric search mode

#### 4.2.1 Observations on 41 upregulated and 28 downregulated miRNAs

To compare our findings with that of the review, we manually explored the papers that the review cited. We found that all the miRNAs, except one, were mentioned in tables, supplementary tables, figures and the whole text beyond the abstracts of papers cited in the review. Since the miRNAs often occur in supplementary tables in these papers and there is no guarantee that the miRNA will occur in the text of the paper, we decided to go for alternate sources of gastric cancer related papers to find the most consistent upregulated and downregulated miRNAs in the context of gastric cancer.

##### Observations on 41 upregulated miRNAs

###### Observation 1: Consistent with review

Out of the 41 upregulated miRNAs, we found 28 miRNAs were upregulated in gastric cancer and stomach cancer.

###### Downregulated miRNA sequences

Contrary to the review’s description, we found 7 out of the 41 miRNAs were consistently downregulated in multiple abstracts. We further analyzed these abstracts and we found that the miRNAs were indeed downregulated. For example, miR-7 was found to be downregulated in PMIDs 22614005 (61), 24573489 (62), 26798443 (63). Similarly, miR-200b was found downregulated in PMIDs 23995857 (64) and 30999814 (65).

###### Closely related upregulated miRNA sequences

From the remaining 6 miRNAs, we found closely related miRNA sequences for 5 of them were upregulated. The review mentioned miR-18b was upregulated (Table S2 in [REF]). However, we found evidence for miR-18a to be upregulated and not for miR-18b. Similarly, we found miR-199a to be upregulated instead of miR-199-5p, miR-301a to be upregulated instead of miR-301, Let-7d and Let-7f to be upregulated instead of Let-7i. We found miR-519 in place of miR-519d in the context of stomach cancer but the expression level was downregulated.

However, miR-1259 was not detected by our tools. Since the review mostly finds upregulated miRNAs from tables or supplementary tables of the papers they survey and our current relation extraction tools are limited to abstracts, we are only able to extract upregulated miRNA mentions from abstracts.

##### Observations on 28 downregulated miRNAs

###### Observation 1: Consistent with review

We found 20 out of 28 downregulated miRNAs mentioned in the review using emiRIT.

###### Upregulated miRNA sequences

From the remaining 8 miRNAs, miR-150 and miR-139 were found to be upregulated.

###### Closely related downregulated miRNA sequences

Instead of miR-320c mentioned in the review, we found miR-320 to be downregulated. Similarly, miR-30b was found to be downregulated instead of miR-30d. However, miR-30d was found to be downregulated in the context of large-intestine cancer and colorectal cancer.

We could not find the remaining 4 miRNAs in the context of stomach cancer or gastric cancer from abstracts of miRNA papers.

#### 4.2.3 Observations on the top 3 upregulated and downregulated miRNAs in the review

The review also stated that from the list of upregulated miRNAs, the most consistently reported miRNAs were miR-21, followed by miR-25, miR-92 and miR-223. From the list of downregulated miRNAs, the authors found that the most consistently reported miRNAs were miR-375, miR-148a followed by miR-638. To find the most consistent upregulated and downregulated miRNAs using emiRIT, we conducted an additional search on the table shown in Figure 12. We filtered this table further to consider only the upregulated or only the downregulated cases and downloaded the resultant tables as a excel separate files. In each file, then looked at the total number of abstracts supporting the regulation (up or down) of a miRNA.

Consistent with the review’s findings, we found miR-21 to be the most reported miRNA upregulated in stomach cancer, extracted from 22 abstracts in emiRIT. The next most frequently reported miRNAs upregulated in abstracts using emiRIT were miR-25 (7 abstracts), miR-27a (7 abstracts), miR-223 (6 abstracts), miR-20a (6 abstracts) and miR-214 (6 abstracts). While miR-92 was not in the top list, we did identify it in 3 abstracts. From the downregulated miRNAs, again consistent with the review’s findings, we found miR-375 to be the most frequently reported miRNA downregulated in stomach cancer, occurring in 9 abstracts in our resource, followed by miR-148a (8 abstracts), miR-218 (6 abstracts) and miR-133b (6 abstracts).

The above case studies showed the amount of information that can be found from the text, specifically abstracts, using emiRIT. The advantage of using our resource is that users get a more comprehensive picture of the miRNA information for their specific requirements.

## 5. Conclusion and Future Work

In this paper, we have described emiRIT, a text-mined based resource for miRNA information. We used different existing and in-house developed text mining tools to capture connections between miRNA-gene, miRNA-disease (cancer), miRNA-biological process and pathways, and miRNA-extracellular locations. Furthermore, instead of just stating an association between a miRNA and a disease, we present a more detailed role of a miRNA in context of disease by distinguishing between i) impact of miRNA on disease process and outcome, ii) influence of miRNA on disease treatment iii) diagnostic role of miRNAs as biomarkers, iv) their role as therapeutic targets in diseases, v) others, where the particular role is not clear, but the miRNA is associated with a disease or regulates a disease.

The output of the different text mining tools is combined, and the output format is unified for easy visualization and navigation of information. All the different miRNA connections are presented to the users via an interface at https://research.bioinformatics.udel.edu/emirit/. Here, the users can easily transition between these connections to get a broader understanding of the role of miRNAs and examine these roles in a context specific to their different information needs. Literature evidence is provided for every result at the abstract and sentence level, which not only increases the confidence of the results extracted from the text but allows users to explore the papers for additional background and experimental context of the results. miRNAs and other bioentities are normalized using standard ontologies to expand querying abilities and integrate with external resources.

We have conducted two case studies to show the extent of miRNA information that can be found through emiRIT. Since the primary function of miRNAs is to regulate gene expression and their dysregulation often leads to diseases, we focused on the target information and differential expressions of miRNAs in the context of diseases for our case studies. We relied on review articles for the case studies to assess the information coverage of our resource because of the unavailability of gold standards that can be used to compare our findings.

In this paper, we attempt to provide an up-to-date and user-friendly resource to facilitate access to comprehensive miRNA information from the literature on a large-scale, enabling users to exploit, interpret and connect existing knowledge to design new investigations and theories. In the future, we plan to extend to diseases other than cancer, and to improve the relation extraction tool capturing the connections between miRNAs and extracellular locations. We also plan to integrate additional miRNA-entity relation knowledge from external databases and provide network visualization through Cytoscape. Finally, we plan to extend our resource to full length PMC open access publications.

