## Supplementary figures and images for "emiRIT: A text-mining based resource for microRNA information"

### Supplemental Figure S2

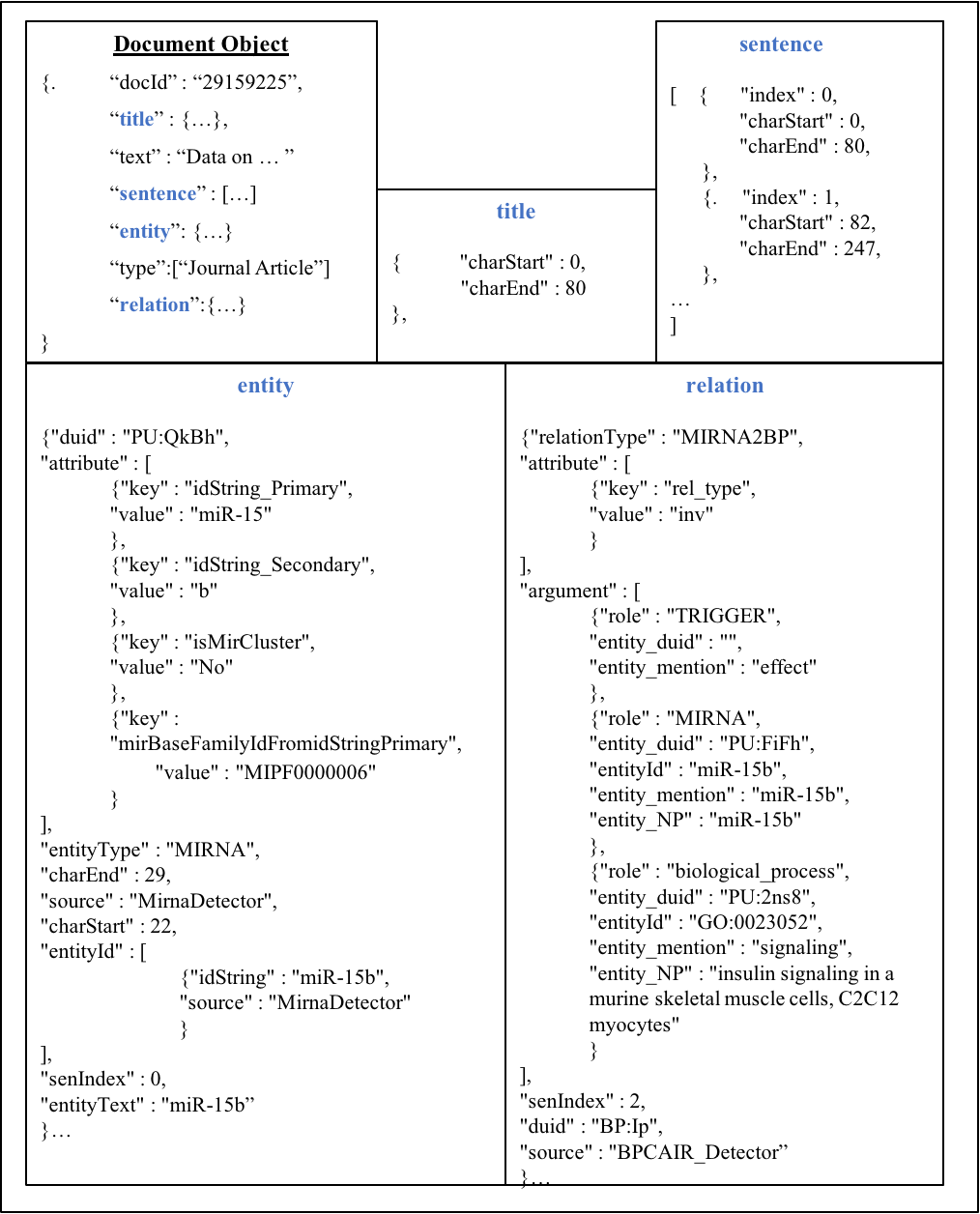

Figure S2: Overview of the JSON format of data stored in the emiRIT database
