## Supplemental Table S1 for "emiRIT: A text-mining based resource for microRNA information"

| Type | Regular Expression | miRNA Example |
| --- | --- | --- |
| 1 | (((?<=[^a-zA-Z])\|^)[a-zA-Z]{1,3}-?)?(microR\|\(miR\)\|miRNA\|micro( \|-)?RNA\|miRN\|miR\|mir)s?(-\|_\|x\| )?([0-9]+([a-g\*p]+[0-9]*)?) | hsa-miR-21,  miR-34a,  microRNA-199-5p,  microRNA-199-3p |
| 2 | (((?<=[^a-zA-Z])\|^)[a-zA-Z]{1,3}-?)?(let\|Let)s?(-\|_\|x\| )?(7[a-g\*p]*)(?=[^0-9]\|$) | Let-7a |
| 3 | (((?<=[^a-zA-Z])\|^)[a-zA-Z]{1,3}-?)?(lin\|Lin)s?(-\|_\|x\| )?(4[a-g\*p]*)(?=[^0-9]\|$) | Lin-4 |

Table S1: Regular expressions to identify base patterns of miRNAs
