## Supplemental Table S3 for "emiRIT: A text-mining based resource for microRNA information"

| # | Upregulated miRNA from review | Upregulated in EmiRIT | Closely related miRNA sequence in EmiRIT | Downregulated in EmiRIT |
| --- | --- | --- | --- | --- |
| s1 | miR‐21 | Y | - | N |
| 2 | miR‐25 | Y | - | N |
| 3 | miR‐92 | Y | - | N |
| 4 | miR‐223 | Y | - | N |
| 5 | miR‐106b | Y | - | N |
| 6 | miR‐106a | Y | - | N |
| 7 | miR‐18a | Y | - | N |
| 8 | miR‐93 | Y | - | N |
| 9 | miR‐17 | Y | - | N |
| 10 | miR‐23a | Y | - | N |
| 11 | miR‐191 | Y | - | N |
| 12 | miR‐19a | Y | - | N |
| 13 | miR‐20a | Y | - | N |
| 14 | miR‐27a | Y | - | N |
| 15 | miR‐214 | Y | - | N |
| 16 | miR‐100 | Y | - | N |
| 17 | miR‐20b | Y | - | N |
| 18 | miR‐425‐5p | Y | - | N |
| 19 | miR‐7 | N | - | Y |
| 20 | miR‐215 | Y | - | N |
| 21 | miR‐135b | Y | - | N |
| 22 | miR‐224 | Y | - | N |
| 23 | miR‐192 | Y | - | N |
| 24 | miR‐221 | Y | - | N |
| 25 | miR‐18b | N | miR-18a (up) | N |
| 26 | miR‐200b | N | - | Y |
| 27 | miR‐194 | Y | - | - |
| 28 | miR‐99b | N | - | Y |
| 29 | miR‐10a | N | - | Y |
| 30 | miR‐15a | N | - | Y |
| 31 | miR‐199‐5p | N | miR-199a (up) | N |
| 32 | miR‐301 | N | miR-301a (up) | N |
| 33 | miR‐519d | N | miR‐519 (down) | N |
| 34 | let‐7i | N | Let 7d (up)  Let 7f (up)  Let-7a (down) | N |
| 35 | miR‐181d | Y | - | - |
| 36 | miR‐185 | Y | - | - |
| 37 | miR‐181a | Y | - | - |
| 38 | miR‐1259 | Not found | - | - |
| 39 | miR‐335 | N | - | Y |
| 40 | miR‐424 | Y | - | - |
| 41 | miR‐542‐3p | N | - | Y |

Table S3: Up or downregulation information found by EmiRIT for upregulated miRNAs from the review Shrestha *et al*., 2014
