## Supplemental Table S4 for "emiRIT: A text-mining based resource for microRNA information"

| # | Downregulated miRNA from review | Downregulated in EmiRIT | Closely related miRNA sequence in EmiRIT | Upregulated in EmiRIT |
| --- | --- | --- | --- | --- |
| 1 | miR‐375 | Y | - | - |
| 2 | miR‐148a | Y | - | - |
| 3 | miR‐30d | N | miR-30b (down) | N |
| 4 | miR‐638 | Y | - | - |
| 5 | miR‐29c | Y | - | - |
| 6 | miR‐155 | Y | - | - |
| 7 | miR‐378 | Y | - | - |
| 8 | miR‐152 | Y | - | - |
| 9 | miR‐30c | Y | - | - |
| 10 | miR‐218 | Y | - | - |
| 11 | miR‐133b | Y | - | - |
| 12 | miR‐150 | N | - | Y |
| 13 | miR‐564 | Not found | - | - |
| 14 | miR‐489 | Y | - | - |
| 15 | miR‐136 | Y | - | - |
| 16 | miR‐197 | Not found | - | - |
| 17 | miR‐923 | Not found | - | - |
| 18 | miR‐490 | Y | - | - |
| 19 | miR‐146a | Y | - | - |
| 20 | miR‐188 | Y | - | - |
| 21 | miR‐34b | Y | - | - |
| 22 | miR‐370 | Y | - | - |
| 23 | miR‐139 | N | - | Y |
| 24 | miR‐513a‐5p | Not found | - | - |
| 25 | miR‐494 | Y | - | - |
| 26 | miR‐320c | N | miR-320 (down) | N |
| 27 | miR‐433 | Y | - | - |
| 28 | miR‐101 | Y | - | - |

Table S4: Up or downregulation information found by EmiRIT for downregulated miRNAs from the review Shrestha *et al*., 2014
